# Adenosine 2B receptor signaling impairs vaccine-mediated protection against pneumococcal infection in young hosts by blunting neutrophil killing of antibody opsonized bacteria

**DOI:** 10.1101/2025.03.18.643957

**Authors:** Shaunna R. Simmons, Alexsandra P. Lenhard, Michael C. Battaglia, Elsa N. Bou Ghanem

## Abstract

**Background/Objective:** Neutrophils are essential for vaccine-mediated protection against pneumococcal infection and impairment in their antibacterial function contributes to reduced vaccine efficacy during aging. However, the signaling pathways controlling neutrophil responses in vaccinated hosts are not fully understood. The extracellular adenosine pathway is a known regulator of neutrophils in naïve hosts. The aim of this study was to test the role of this pathway in neutrophil function and protection against infection upon vaccination across host age.

**Methods:** To test the role of adenosine in the antimicrobial activity of neutrophils against antibody-opsonized pneumococci, we used bone marrow derived neutrophils isolated from wild type or specific adenosine receptors knock-out mice. To measure the effect of adenosine receptor signaling *in vivo*, we treated vaccinated mice with agonists or antagonists specific to the different adenosine receptors prior to pulmonary challenge with pneumococci and assessed bacterial burden and clinical score post infection.

**Results:** We found that signaling via the adenosine 2B (A2BR) but not A2A or A1 receptor diminished intracellular pneumococcal killing following antibody-mediated uptake in young hosts. *In vivo*, agonism of A2BR significantly worsened pneumococcal infection outcome in young, vaccinated mice. In contrast, A2BR signaling had no effect on intracellular bacterial killing by neutrophils from aged mice. Further, *in vivo* A2BR inhibition had no effect on pneumococcal disease progression in aged, vaccinated mice.

**Conclusions:** A2BR signaling reduced pneumococcal vaccine-mediated protection by impairing neutrophil antimicrobial activity against antibody-opsonized bacteria in young hosts. However, inhibiting this pathway was not sufficient to boost responses in aged hosts.

## INTRODUCTION

*Streptococcus pneumoniae* (pneumococcus) are Gram positive, encapsulated bacteria that are the leading cause of bacterial pneumonia globally, and the leading cause of community acquired bacterial pneumonia in those over the age of 65 [1–4]. Older adults are at increased risk of contracting pneumococcal disease, with an increased number of cases and increased mortality rate occurring in those over the age of 65 [5]. Increased risk of pneumococcal pneumonia persists in this population despite the availability of vaccines [5]. Licensed pneumococcal vaccines include the pneumococcal conjugate vaccine (PCV) that is currently recommended for older adults [6]. These vaccines target the bacterial polysaccharides but have reduced efficacy in protection against pneumococcal pneumonia in older adults [7–9]. Reduced vaccine efficacy with age can be partly attributed to immunosenescence, the age-related decline in immune system function [9]. Vaccination helps mediate pneumococcal bacterial clearance in a variety of ways; antibody binding on the surface of bacteria prevents bacteria from binding to lung epithelium and can induce complement-mediated killing [10]. Additionally, opsonization of pneumococcus by antibodies promotes uptake and clearance by phagocytes, including neutrophils also known as polymorphonuclear leukocytes (PMNs) [9, 10]. However, both anti-pneumococcal antibody and neutrophil functionality decline with host aging [9, 11, 12].

PMNs are important mediators of pneumococcal disease outcome and are one of the first cells recruited to the lung following pneumococcal pneumonia [13, 14]. PMNs kill bacteria in a variety of ways including phagocytosis and subsequent intracellular bacterial killing, degranulation and release of preformed antimicrobial products, production of reactive oxygen species (ROS) and through release of neutrophil extracellular traps (NETs) [15]. PMNs are essential for host defense against pneumonia; neutropenic patients are at higher risk of contracting pneumonia and in mouse models of pneumococcal infection, depletion of PMNs prior to pulmonary challenge resulted in significantly higher bacterial burden and mortality [1, 11, 14, 16, 17]. We also showed that PMNs are important for protection of vaccinated hosts, where depletion of PMNs in PCV immunized mice resulted in a significant reduction in host survival upon pulmonary challenge with *S. pneumoniae* [11]. We found that PMN antimicrobial activity against antibody opsonized bacteria declined in vaccinated aged hosts and that this decline persisted even when antibodies were isolated from young controls, suggesting an intrinsic defect in PMN function [11]. We specifically found that following antibody-mediated uptake there was a significant decline in the ability of PMNs isolated from aged mice to kill *S. pneumoniae* intracellularly. Adoptive transfer of PMNs from young, naïve mice into aged, vaccinated mice boosted host protection *in vivo* suggesting that boosting PMN function in aged, vaccinated hosts can improve pneumococcal disease outcome [11].

A known regulator of immune cell function is the extracellular adenosine (EAD) pathway. During infection and inflammation ATP leaks from damaged cells and is converted to extracellular adenosine by dephosphorylation by the extracellular enzymes CD39 and CD73 [18, 19]. EAD can then act on the four G-protein coupled adenosine receptors, A1, A2A, A2B, and A3 that can have opposing downstream effects on immune cell responses [18, 19]. Adenosine receptors are expressed on PMNs and this pathway has been shown to regulate PMN responses to bacterial infection, or upon stimulus by bacterial products [19]. For example, A1 receptor (A1R) signaling was necessary for efficient PMN influx to the lungs following pneumococcal infection [20] and was also required for efficient intracellular killing of *S. pneumoniae* by PMNs [21]. Activation of the A2A receptor (A2AR) was shown to decrease MMP-9 secretion by fMLP activated PMNs [22, 23]. A 2B receptor (A2BR) knock-out mice had improved survival upon pulmonary infection with *Klebsiella pneumoniae* and A2BR^-/-^ PMNs had higher production of NETs and increased bactericidal activity compared to wild type PMNs [24]. Similarly, A2BR^-/-^ mice were more resistant to pneumococcal infection and A2BR^-/-^ PMNs had increased mitochondrial ROS production that resulted in increased bactericidal activity against *S. pneumoniae* [25, 26].

Although EAD has been shown to regulate PMN response in naïve hosts, our understanding of how EAD controls PMN antimicrobial response in immune hosts remains limited. The way PMNs interact with bacteria in naïve versus vaccinated hosts can significantly differ due to the differences in opsonins. The pneumococcal capsule helps bacteria evade phagocytosis [4], however, pneumococci can be opsonized by complement or antibody deposition on the bacterial surface, which helps promote phagocytosis and clearance of the bacteria via activation of complement receptors (CR) and Fc receptors on the neutrophil surface [4, 27, 28]. It is important to understand the differences in complement-mediated and antibody-mediated responses by PMNs because activation of theses receptors results in distinct signaling pathway activation in PMNs [27, 29, 30]. It was shown that phagocytosis of complement coated, or IgG coated beads by PMNs induced differing levels of ROS, differences in phagocytosis and resulted in receptor specific changes in gene expression [31]. To determine the pathways controlling PMN responses in a vaccinated host, we asked if EAD signaling regulates killing of *S. pneumoniae* following antibody mediated uptake by PMNs and if this pathway can be targeted to reverse the decline in vaccine protection in aged hosts.

## MATERIALS AND METHODS

### Mice

Young (2-3 months) and old (20-22 months) C57BL/6 (B6) male mice were purchased from Jackson Laboratories and from the National Institute on Aging. A2BR^-/-^ on a B6 background (B6.129P2-Adora2btm1Till/J) and A2AR^-/-^ mice on a Balb/c background (C;129S-Adora2atm1Jfc/J) were purchased from Jackson Laboratory and bred at our facility. Age matched wild type (WT) C57BL/6J and Balb/cJ were used as controls. All mice were housed in specific pathogen free housing for at least two weeks prior to use in experiments. A2BR^-/-^ mice bred at our facility were aged for 18 months.

### Ethics statement

All animal studies were performed in accordance with the recommendations in the Guide for the Care and Use of Laboratory Animals and in accordance with the University at Buffalo Institutional Animal Care and Use Committee guidelines.

### Bacteria

#### Streptococcus pneumoniae

TIGR4 strain (serotype 4), wild type (WT) and pneumolysin deletion mutant (*Δply*) were a gift from Andrew Camilli [32]. Bacteria were grown to mid-exponential phase at 37⁰C at 5% CO_2_ in Todd Hewitt broth supplemented with 0.5% yeast extract and oxyrase as previously described [33].

### PMN Isolation

Bone marrow (BM) was isolated from the femurs and tibias of naïve, uninfected mice by cutting the end of each bone, flushing with RPMI supplemented with 10% FBS and 2mM EDTA, and strained via 100μm cell strainers. Red blood cells were lysed, and remaining cells were washed and resuspended in PBS. PMNs were separated out by density centrifugation using Histopaque 1119 and 1077 as previously described [34]. PMNs were then resuspended at the indicated concentrations in Hank’s Balanced Salt Solution/0.1% gelatin with no Ca^2+^ or Mg^2+^ and kept on ice until use. The purity of PMNs was confirmed using flow cytometry and 85-90% of enriched cells were positive for Ly6G and CD11b.

### Generation of Immune Sera

Young WT mice were vaccinated intramuscularly into the caudal thigh muscle with the pneumococcal conjugate vaccine (PCV) Prevnar-13^®^ (Wyeth pharmaceuticals). Four weeks following vaccination, mice were euthanized, and blood was collected via portal vein snip and centrifuged at 9000 rpm to separate the sera. Sera were then stored at −80⁰C until use. To heat inactivate (HI) the complement proteins in the immune sera prior to use, sera were incubated at 56⁰C for 40 minutes as previously described [35]. We previously demonstrated that heat inactivation prevented complement deposition but had no effect on anti-capsular IgG binding to bacterial surfaces [11].

### Mouse Infection and Treatment

Mice were vaccinated with Prevnar-13^®^ (PCV-13) four weeks prior to treatment and infection. Mice were infected with 50ul of *S. pneumoniae* TIGR4 at concentrations indicated via assisted aspiration, which we refer to here as intratracheally (i.t.). Briefly, mice were anesthetized using isoflurane and pneumococcus delivered to the lungs by pipetting bacteria into the trachea with the tongue gently pulled to the side. For treatment with specific adenosine receptor agonists and antagonists, mice were injected intraperitoneally (i.p) 18 hours prior to, at the time of, and 18 hours post infection. Young mice were treated with specific A2B receptor agonist Bay 60-6583 dissolved in DMSO, filter sterilized by passing through a 0.22μm filter prior to use and given to mice via intraperitoneal injection at a concentration of 2mg/kg. Old mice were treated with specific A2B receptor antagonist MRS 1754 dissolved in DMSO, filter sterilized by passing through a 0.22μm filter prior to use and given to mice via intraperitoneal injection at a concentration of 2.5mg/kg. Where indicated, old mice were also treated with specific A2A receptor antagonist 3,7-Dimethyl-1-propargylxanthine dissolved in DMSO, filter sterilized by passing through a 0.22μm filter prior to use and given to mice via intraperitoneal injection at a concentration of 5mg/kg simultaneously with treatment of A2BR antagonist MRS 1754. All control mice were treated with diluted DMSO as vehicle control. 24 hours post infection, each mouse was assessed for clinical signs of disease (clinical score) and subsequently euthanized and blood collected via portal vein snip to determine bacteremia. After blood collection, mice were perfused with sterile PBS through the right ventricle and lungs and brain harvested. Following harvest, lung and brain samples were homogenized in sterile PBS, diluted in PBS, and plated on blood agar to enumerate colony forming units (CFU).

### Gentamicin Protection Assay

PMNs were isolated from the bone marrow of unvaccinated mice as indicated as we previously found that vaccination did not alter intrinsic PMN antimicrobial activity against *S. pneumoniae* [11]. PMNs were then infected at a MOI of 25 with *Δply S. pneumoniae* TIGR4, pre-opsonized with 3% naïve or heat inactivated immune sera for 10 minutes at 37°C. Gentamicin (100μg/ml) was then added for 30 minutes to kill remaining extracellular bacteria. PMNs were then washed and resuspended in HBSS/0.1% gelatin. Half of this resuspension was diluted and immediately plated on blood agar plates to determine the amounts of engulfed bacteria. The other half of PMNs were incubated for an additional 15 minutes at 37°C and then plated on blood agar to enumerate remaining viable bacteria. The percentage of engulfed bacteria (at 10 minutes) that was killed was then calculated. We previously found no significant differences between PMNs overall killing of wild-type and *Δply S. pneumoniae* TIGR4 but that presence of the pore forming toxin pneumolysin (PLY) prevented detection of intracellular bacteria using gentamicin protection as the pores allowed influx of the antibiotic into the PMNs [33]. Where indicated, PMNs were pretreated for 30 minutes at 37°C with specific A1 agonist, 2-Chloro-N6-cyclopentyladenosine at a concentration of 2Ki (2nM) prior to infection.

### CRAMP ELISA

PMNs were infected with *S. pneumoniae* TIGR4 pre-opsonized with 3% naïve or heat inactivated immune sera at a MOI of 2. Control wells were mock infected with PMNs alone. PMNs were infected for 40 minutes at 37⁰C. Cells were then centrifuged to separate the supernatants and pellets. Pellets were lysed with RIPA lysis buffer with 0.1% Tx-100 and analyzed for Cathelicidin antimicrobial peptide (CRAMP ELISA kit, mybiosource) as per manufacturer’s instructions.

### Mitochondrial ROS

PMNs isolated from the BM were infected with *S. pneumoniae* TIGR4 pre-opsonized with 3% heat inactivated immune sera at a MOI of 10. Control wells were mock infected with PMNs and sera alone. MitoSOX^TM^ Red mitochondrial superoxide indicator for live cell imaging (Invitrogen) was then added and the 96 well plate was immediately placed in a plate reader to detect mitochondrial ROS production. Readings were taken every minute over a 1-hour time period, excitation 485nm and emission 585nm.

### Statistics

Statistics were analyzed using Prism 9 (GraphPad). Graphs are shown as Mean +/- standard deviation with each data point representing a different mouse. Normality of data was tested via Shapiro-Wilk test before statistical analysis was performed. Significant difference between groups was determined by One Sample t and Wilcoxon test, and Mann-Whitney test as indicated. *p <0.05* was considered significantly different and is indicated in figures with *.

## RESULTS

### A1 receptor signaling activation does not enhance intracellular killing by PMNs from aged mice following antibody-mediated uptake

Previously published work from our lab showed that signaling through the A1 adenosine receptor was necessary for efficient intracellular bacterial killing by PMNs from young and old, naïve mice [21]. To test the role of this receptor in intracellular killing following antibody-mediated bacterial uptake, we performed a gentamicin protection assay using PMNs isolated from the BM of aged mice. PMNs were treated with a specific A1 receptor agonist (2-Chloro-N6-cyclopentyladenosine) or vehicle control (VC) and infected with *S. pneumoniae Δply* TIGR4 opsonized with heat inactivated (HI) immune sera or naïve sera as a control. To test intrinsic PMN function independent of the confounding effects of the age-driven decline in antibody responses, sera were collected from young controls. Consistent with prior reports [21], we found that when *S. pneumoniae* was opsonized with sera from naïve mice, treatment of PMNs with A1 agonist significantly increased the ability of these PMNs to kill intracellular pneumococcus when compared to vehicle control (**Figure 1**). However, when *S. pneumoniae* was opsonized with HI immune sera, A1 agonism had no effect on the ability of PMNs from old mice to kill these bacteria intracellularly (**Figure 1**). These data show that unlike in naïve old hosts, the A1 adenosine receptor is not regulating intracellular killing by PMNs following antibody-mediated bacterial uptake in old hosts.

**Figure 1:**
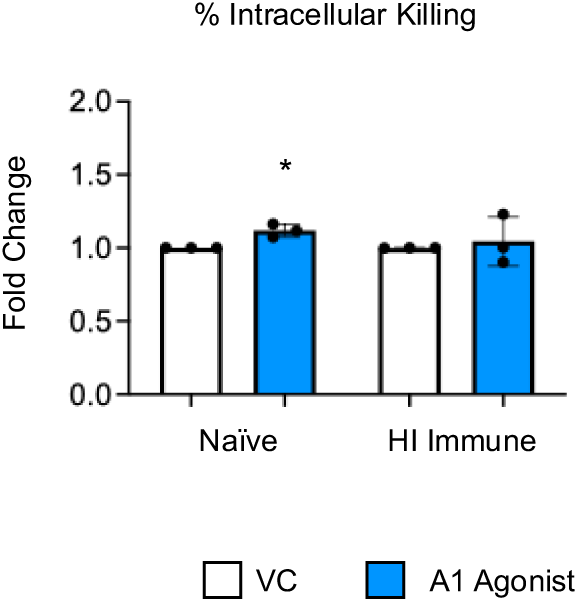
A1 receptor does not control intracellular bacterial killing following antibody-mediated uptake by PMNs from aged mice. PMNs were isolated from the BM of C57BL/6 old (20-22 months) mice and pretreated with A1 agonist or vehicle control (VC). PMNs were then infected with *S. pneumoniae Δply* TIGR4, pre-opsonized with naïve or HI immune sera. Gentamicin was added to kill extracellular bacteria not taken in. Reactions were plated to on blood agar to enumerate CFU. The % of engulfed bacteria that was killed was then calculated. Fold change in bacterial killing of A1 agonist versus VC treated cells are shown. Data from four separate experiments with n=4 biological replicates are pooled. * denotes significantly different from one (*p<0.05)* as determined by One Sample t and Wilcoxon test.

### A2B receptor signaling is detrimental to intracellular bacterial killing by PMNs following antibody-mediated uptake

To test if other adenosine receptors regulate intracellular bacterial killing following antibody-mediated uptake, we repeated the gentamicin protection assay using PMNs isolated from the bone marrow of young, A2AR^-/-^ mice and their wild type (WT) controls. As A2AR was previously found to be required for antibody responses to pneumococcal polysaccharides [36], to test intrinsic PMN function, sera were collected from WT controls. We found that there was no difference between knock out or wild type PMNs in intracellular killing of *S. pneumoniae* when these bacteria were opsonized with naïve, or heat inactivated immune sera indicating that the A2A receptor is not regulating intracellular bacterial killing (**Figure 2**). We next performed this assay using PMNs isolated from the bone marrow of young A2BR^-/-^ mice and their WT controls (**Figure 3**). To test intrinsic PMN function, sera were again collected from WT controls. We found that in PMNs isolated from young mice, when *S. pneumoniae* is opsonized with HI immune sera, intracellular killing of these bacteria significantly increased in the absence of A2B adenosine receptor (**Figure 3**), indicating that signaling through this receptor is detrimental to bacterial killing. A2B receptor signaling did not have an effect on intracellular killing when *S. pneumoniae* were opsonized with sera from naïve mice (**Figure 3**). These data indicate a specific role for A2B receptor signaling in intracellular killing of pneumococcus following antibody-mediated uptake but not complement-mediated uptake in young hosts.

**Figure 2:**
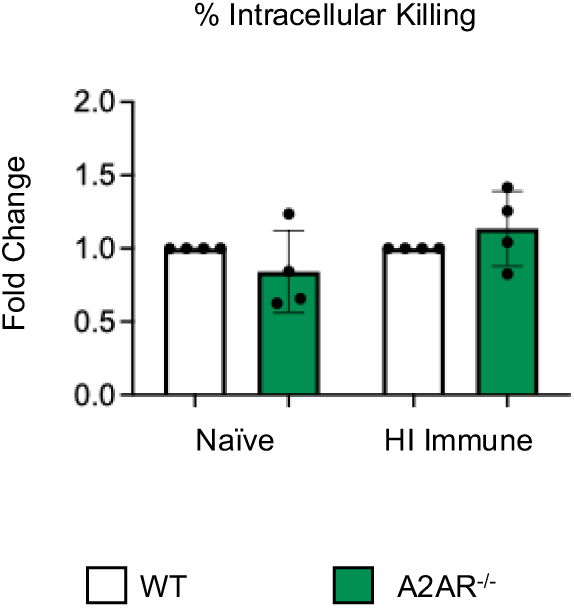
A2A receptor shows no role in controlling intracellular bacterial killing following antibody-mediated uptake by PMNs. PMNs were isolated from the bone marrow of young (2-3 months) wild type (WT) BALB/c, or A2AR^-/-^ mice. PMNs were infected with *S. pneumoniae Δply* TIGR4, pre-opsonized with naïve or HI immune sera. Gentamicin was added to kill extracellular bacteria not taken in. Reactions were plated to on blood agar to enumerate CFU. The % of engulfed bacteria that was killed was then calculated. Fold change in bacterial killing of A2AR^-/-^ versus WT PMNs are shown. Data from four separate experiments with n=4 biological replicates are pooled.

**Figure 3:**
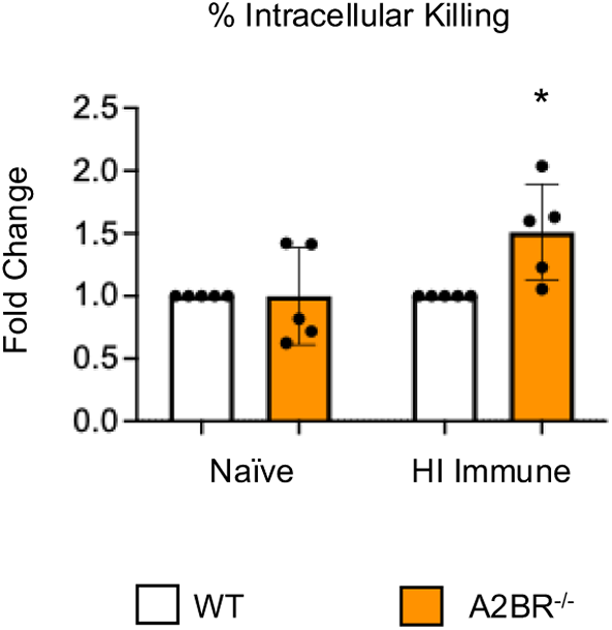
The absence of A2B receptor increases intracellular killing by PMNs following antibody-mediated uptake. PMNs were isolated from the bone marrow of young wild type (WT) C57BL/6, or A2BR^-/-^ mice. PMNs were infected with *S. pneumoniae Δply* TIGR4, pre-opsonized with naïve or HI immune sera. Gentamicin was added to kill extracellular bacteria not taken in. Reactions were plated on blood agar to enumerate CFU. The % of engulfed bacteria that was killed was then calculated. Fold change in bacterial killing of A2BR^-/-^ versus WT PMNs are shown. Data from five separate experiments with n=5 biological replicates are pooled. * denotes significantly different from one (*p<0.05)* as determined by One Sample t and Wilcoxon test.

### PMNs from A2BR^-/-^ mice upregulate intracellular levels of CRAMP following *S. pneumoniae* infection

To begin to understand how PMNs isolated from A2BR^-/-^ mice had improved intracellular killing of pneumococcus after antibody-mediated uptake, we first analyzed mitochondrial ROS production. Mitochondrial ROS is necessary for pneumococcal killing and was shown to be elevated in naïve A2BR^-/-^ mice [26]. We performed an assay to quantify ROS produced by the mitochondria using MitoSOX dye. We found that following infection with *S. pneumoniae* pre-opsonized with HI immune sera, PMNs from both WT and A2BR^-/-^ mice upregulated mitochondrial ROS production to the same extent within the first twenty minutes but that the response was better sustained over time in WT PMNs (**Figure 4A and B**). These data suggest that mitochondrial ROS production does not explain the increased intracellular killing we observe in A2BR^-/-^ PMNs (**Figure 3**). We next analyzed intracellular granule components. PMNs kill intracellular bacteria through several mechanisms, including delivery of antimicrobial peptides and enzymes to pneumococcus contained in phagosomes [37, 38]. Previously, we found that a decline in intracellular killing of antibody opsonized *S. pneumoniae* by PMNs from aged mice when compared to young controls was accompanied by a decline in intracellular cathelicidin-related antimicrobial peptide (CRAMP) [11]. To test if A2B receptor signaling was regulating intracellular CRAMP levels, we performed CRAMP ELISA on cell pellets collected from WT and A2BR^-/-^ PMNs infected with *S. pneumoniae* opsonized with HI immune sera (**Figure 4C and D**). We found that upon infection, there was twice as much intracellular CRAMP in PMNs from A2BR^-/-^ mice when compared to WT controls (**Figure 4C**). When compared to uninfected baseline, PMNs from A2BR^-/-^ mice significantly upregulated CRAMP levels upon infection (**Figure 4D**). These data suggest that A2BR signaling inhibits upregulation of the anti-pneumococcal peptide CRAMP in response to infection with antibody opsonized bacteria.

**Figure 4:**
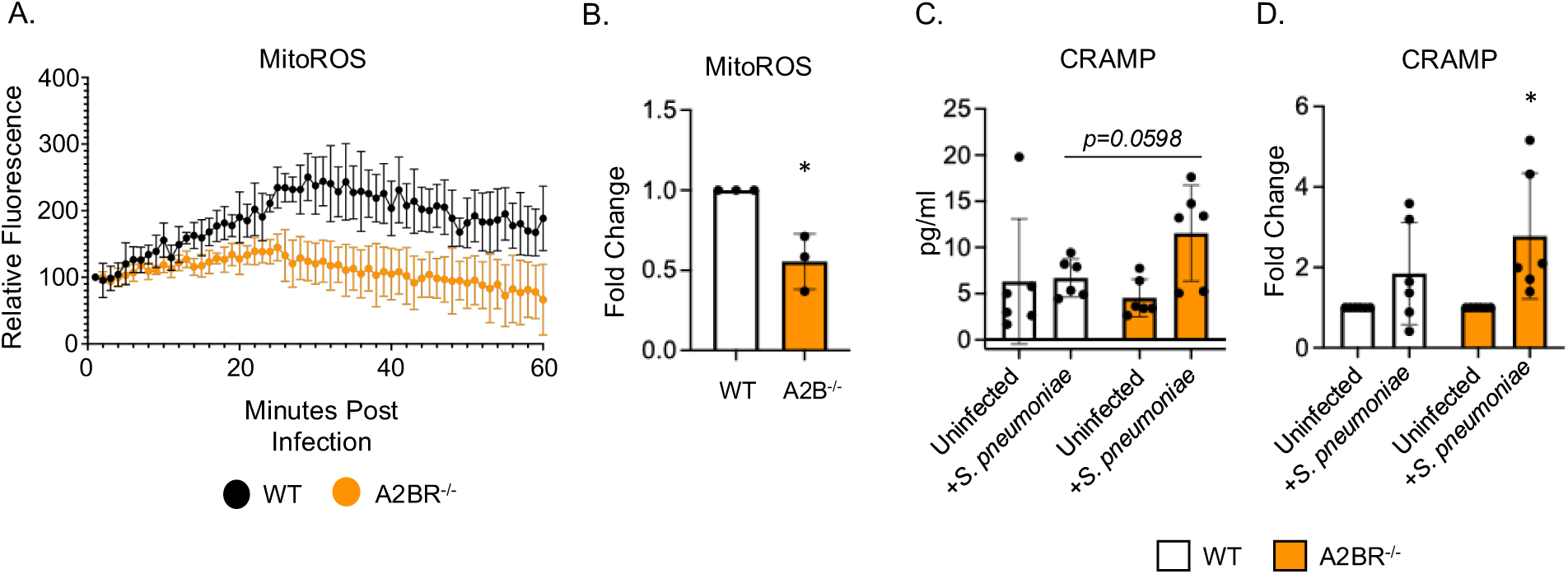
A2BR signaling blunts intracellular CRAMP levels in PMNs following antibody-mediated bacterial uptake. PMNs were isolated from the bone marrow of young wild type C57BL/6, or A2BR^-/-^ mice. PMNs were infected with *S. pneumoniae* opsonized with HI immune sera. (A and B) Mitochondrial ROS production was determined uisng MitoSOX. (A) Mitochondrial ROS production over the course of 1 hour following infection. Representative data of 1 of 3 separate experiments are shown. (B) Fold changes in mitochondrial ROS production by A2BR^-/-^ PMNs from wild type controls were calculated using area under the curve. Data are pooled from separate experiments and * indicates significantly different from 1 as determined by One Sample t and Wilcoxon test. (C and D) Cell pellets were collected and lysed and CRAMP ELISA performed. Pooled data from separate mice (C) and corresponding fold changes from uninfected baselines (D) are shown. (C) number indicates the *p* value as determined by unpaired student’s t test and (D) * indicates significantly different from 1 as determined by One sample t and Wilcoxon test.

### A2BR agonism *in vivo* worsens pneumococcal disease outcome in vaccinated, young mice

Since PMNs from A2BR^-/-^ PMNs had improved intracellular bacterial killing following antibody-mediated uptake compared to WT controls (**Figure 3**), we next wanted to test if A2B receptor signaling was important during *in vivo* infection in a vaccinated host. To test this, young A2BR^-/-^ and WT mice were vaccinated with PCV and 4 weeks later infected with *S. pneumoniae* TIGR4 and monitored for bacteremia, clinical score, and survival for 7 days post infection (**Supplemental Figure 1**). We found that when vaccinated, both WT and A2BR^-/-^ mice were fully protected from pneumococcal disease following infection. We found no incidence of bacteremia in either WT or A2BR^-/-^ mice (**Supplemental Figure 1A**), no differences in clinical score (**Supplemental Figure 1B**), and 100% of all infected mice survived the infection (**Supplemental Figure 1C**). As even WT controls were fully protected and therefore improved protection in A2BR^-/-^ mice was not possible to observe, we instead asked if activation of the A2B receptor impairs protection of vaccinated hosts upon *in vivo* infection. To test this, young WT mice were vaccinated with PCV and 4 weeks later treated with specific A2BR agonist (Bay 60-6583) or VC 18 hours before, at the time of, and 18 hours post infection with *S. pneumoniae*. Clinical score and CFU were then analyzed 24 hours post infection. Following infection, the group that received the A2BR agonist had significantly higher clinical score (**Figure 5A**) and significantly higher bacterial burden in the lungs (**Figure 5B**). These data show that during pneumococcal infection, activation of A2BR worsens disease outcome in young, vaccinated hosts.

**Figure 5:**
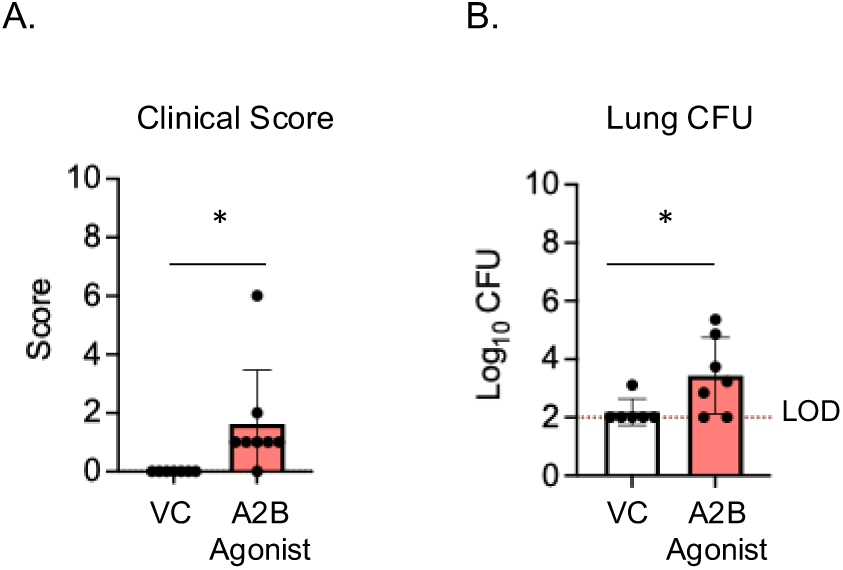
A2BR activation *in vivo* worsens disease outcome in young, vaccinated mice. Young C57BL/6 mice were vaccinated with PCV and 4 weeks later infected with 5×10^6^ CFU *S. pneumoniae* TIGR4. 18 hours prior to, at the time of, and 18 hours post infection mice were treated i.p with a specific A2BR agonist or VC. 24hpi mice were assessed for clinical signs of disease (A). Organs were harvested 24hpi and CFU in the lungs (B) were enumerated by plating on blood agar. * indicates significant differences (*p<0.05)* as determined by Mann-Whitney test. The dashed line indicates the limit of detection (LOD). Data are pooled from two separate experiments and each data point represents an individual mouse.

### Absence of A2BR signaling does not boost intracellular killing following antibody-mediated uptake in aged mice

Previously, we found that PMNs from aged mice have a significant defect in intracellular killing of *S. pneumoniae* following antibody-mediated uptake when compared to PMNs isolated from young mice and that resulted in subpar protection following vaccination [11]. To determine if the A2B receptor can be targeted to improve bacterial killing by PMNs from aged hosts, A2BR^-/-^ mice were aged for 18 months. PMNs were isolated from the bone marrow and gentamicin protection assay used to determine intracellular killing by WT and A2BR^-/-^ PMNs infected with *S. pneumoniae* opsonized with naïve, or HI immune sera. To test intrinsic PMN function, sera were collected from young WT controls. There was no difference in intracellular killing between WT and A2BR^-/-^ PMNs from aged mice in either of the conditions tested (**Figure 6**). These data suggest that removal of A2B receptor is not sufficient to boost intracellular bacterial killing following antibody-mediated uptake by PMNs from aged mice.

**Figure 6:**
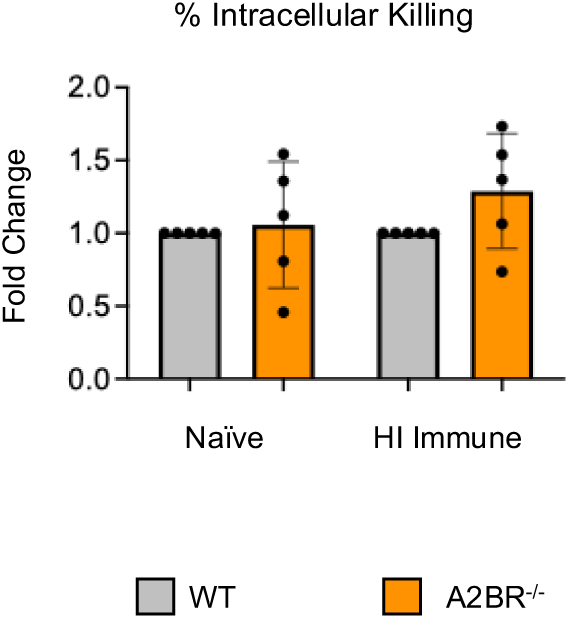
The absence of A2B receptor signaling does not boost intracellular killing by PMNs from aged mice. PMNs were isolated from the bone marrow of old (18 months) wild type (WT) C57BL/6, or A2BR^-/-^ mice. PMNs were infected with *S. pneumoniae Δply* TIGR4, pre-opsonized with naïve or HI immune sera. Gentamicin was added to kill extracellular bacteria not taken in. Reactions were plated to on blood agar to enumerate CFU. The % of engulfed bacteria that was killed was then calculated. Fold change in bacterial killing of A2BR^-/-^ versus WT PMNs are shown. Data from five separate experiments with n=5 biological replicates are pooled.

### Inhibition of A2BR signaling *in vivo* does not improve protection following pneumococcal infection in aged, vaccinated hosts

As activating A2B receptor signaling in young, vaccinated mice worsened disease outcome, we next asked if A2BR inhibition *in vivo* in old, vaccinated mice would improve host resistance to infection. Aged WT mice were vaccinated with PCV and 4 weeks later treated with specific A2B receptor inhibitor (MRS 1754) or VC 18 hours before, at the time of, and 18 hours post infection with *S. pneumoniae*. MRS 1754 was used at the same dose and treatment regimen that we had previously found to improve resistance of young naïve mice to infection (data not shown). Clinical score and CFU were then analyzed 24 hours post infection. When compared to young VC treated mice, old VC treated mice had higher clinical score and bacterial burden in the lungs and blood (**Figures 3B and 3C and Figures 7B and 7C**). These data confirmed that aged hosts are not protected from pneumococcal infection despite vaccination as we had previously reported [11]. When comparing old VC treated to A2BR inhibitor treated mice there was no significant difference in clinical score (**Figure 7A**), bacterial burden in the lungs (**Figure 7B**), blood (**Figure 7C**), or brain (**Figure 7D**). These data show that inhibition of A2B receptor signaling in vaccinated, aged hosts does not reverse the age-related susceptibility to pneumococcal infection. As both A2A and A2B receptors are low affinity adenosine receptors that are both G_s_ coupled and have high homology [19, 39], we wanted to test if signaling through the A2A receptor was compensating *in vivo* upon A2BR inhibition in vaccinated aged mice. Aged WT mice were vaccinated and treated with both specific A2B receptor antagonist (MRS 1754) and specific A2A receptor antagonist (3,7-Dimethyl-1-propargylxanthine) 18 hours prior to, at the time of, and 18 hours post infection. At 24 hours post infection, clinical score and CFU were analyzed. When both A2AR and A2BR were inhibited, there was still no difference from VC treated or A2BR inhibition alone in clinical score (**Figure 7A**), bacterial burden in the lungs (**Figure 7B**), blood (**Figure 7C**), or brain (**Figure 7D**). These data suggest that combined inhibition of A2BR and A2AR in vaccinated hosts does not reverse the age-related susceptibility to pneumococcal infection.

**Figure 7:**
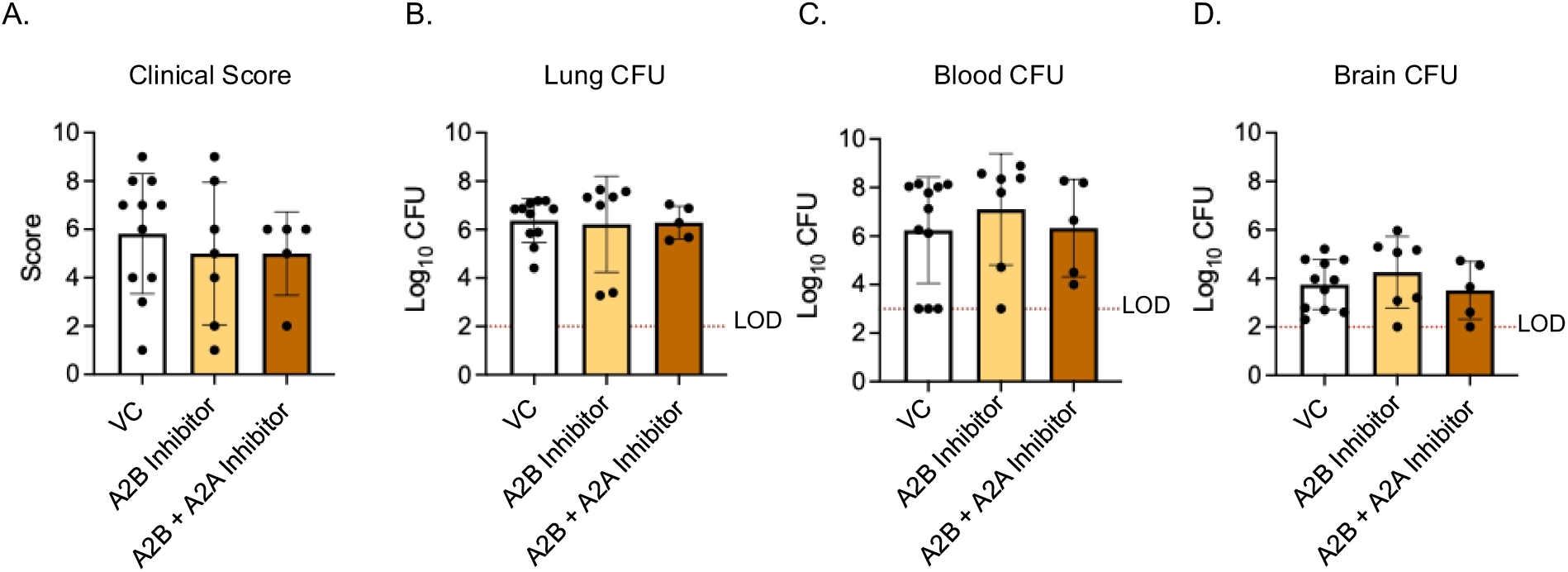
A2BR inhibition *in vivo* has no effect on disease outcome in aged, vaccinated mice. Old (20-22 months) C57BL/6 mice were vaccinated with PCV and 4 weeks later infected with 5×10^6^ CFU *S. pnuemoniae* TIGR4. 18 hours prior to, at the time of, and 18 hours post infection mice were treated i.p with either a specific A2BR inhibitor alone, a combination of specific A2BR inhibitor and A2AR inhibitor or VC. 24hpi mice were assessed for clinical signs of disease (A). Organs were harvested 24hpi and CFU in the lungs (B), blood (C), and brain (D) were enumerated by plating on blood agar. The dashed line indicates the limit of detection (LOD). Data are pooled from two separate experiments and each data point represents an individual mouse.

## DISCUSSION

Extracellular adenosine signaling is a known regulator of PMN responses to infection and is necessary for host protection against pneumococcal infection. Activation of each of the four adenosine receptors has differing effects on PMN responses. A1 receptor activation has been found to be stimulatory, promoting PMN recruitment to the lungs, adhesion, intracellular killing, and superoxide production [19–21, 40]. Conversely, A2B receptor activation has been found to be inhibitory, reducing bactericidal activity, and inhibiting both PMN recruitment and superoxide production [19, 24, 26, 40]. However, the role of these receptors in immune hosts is not well known. In this study we found that unlike in naïve models of infection, A1 receptor signaling does not boost intracellular killing of *S. pneumoniae* following antibody-mediated uptake by PMNs from aged hosts. In PMNs isolated from young hosts, A2B but not A2A receptor signaling had a role with A2BR signaling being detrimental to intracellular killing of pneumococcus following antibody-mediated uptake. However, we found that removal or inhibition of A2BR was not sufficient to correct the age-driven defects in PMN antimicrobial activity against antibody bound pneumococci or the age-driven decline in pneumococcal vaccine efficacy. These data indicate that the EAD pathway components controlling PMN responses in naive and vaccinated hosts are different and distinct and that changes in host aging also effects this pathway.

A1R and A2BR signal differently when activated. A1 is a high affinity adenosine receptor meaning lower levels of adenosine are needed to activate it [41, 42]. A1R is a G_i/o_ coupled GPCR its activation inhibits adenylyl cyclase and inhibits cAMP production [41]. A2BR is a low affinity adenosine receptor, requiring higher μM levels of adenosine to activate signaling and is a G_s_ coupled GPCR [39, 41, 42]. Activation of this receptor stimulates adenylyl cyclase and stimulates cAMP production [41]. Therefore, activation of A1R and A2BR can have opposing downstream signaling effects resulting in differences in PMN responses. In prior work from our lab, we found that when *S. pneumoniae* is opsonized with sera from a naïve, unvaccinated mouse, A1R signaling is required for PMN bacterial killing [21]. Specifically, intracellular killing following complement-mediated uptake was impaired when A1R was inhibited in PMNs from young mice and agonism of this receptor in PMNs isolated from aged mice enhanced intracellular killing [21]. Here, we confirmed that agonism of A1R in PMNs from aged mice increased intracellular killing following complement-mediated uptake. However, following antibody-mediated uptake, A1R agonism had no effect on intracellular killing of pneumococcus in aged hosts. This suggests that A1R signaling is required for complement-mediated but not antibody-mediated PMN response. We found in PMNs isolated from young mice that in the absence of A2BR signaling intracellular killing following antibody mediated uptake increased significantly, indicating that A2BR is detrimental to intracellular killing of antibody opsonized pneumococcus. Previously our lab found that A2BR signaling impairs PMN antimicrobial activity in naive hosts where agonism of this receptor impaired PMN killing of *S. pneumoniae* opsonized with complement while PMNs from A2BR^-/-^ mice had increased killing [26]. These data indicate that unlike A1R signaling A2BR signaling is inhibitory of PMN antimicrobial responses to *S. pneumoniae* is independent of opsonin and that it impairs both complement-mediated and antibody-mediated PMN responses.

PMNs kill *S. pneumoniae* intracellularly through fusion of preformed antimicrobial granules with the phagosome and through intracellular ROS production [37, 43]. In *S. pneumoniae* infection, ROS produced by the NADPH oxidase complex is not necessary for control of this pathogen, however, ROS produced by the mitochondria was required for pneumococcal killing [26]. Previously, it was found that PMNs from A2BR^-/-^ mice had increased mitochondrial ROS production and enhanced overall killing of *S. pneumoniae* opsonized with sera from naïve hosts [26]. Here we found that PMNs from young A2BR^-/-^ mice kill *S. pneumoniae* more efficiently intracellularly following antibody-mediated uptake, however that was not due higher levels of mitochondrial ROS production. While A2BR is detrimental to bacterial control in both naïve and vaccinated models of infection, these data indicate that the mechanism of PMN antimicrobial responses between complement and antibody activation differ. This could be attributed to differing signaling pathways that are activated when complement receptors or Fcγ receptors are activated to initiate phagocytosis. PMNs generate ROS through activation of the NADPH oxidase complex by activation of GPCRs, complement and Fc receptor signaling [43]. However, less is known about signaling pathways that generate Mitochondrial ROS in PMNs. In macrophages, mitochondrial ROS is generated through activation of toll like receptors, proinflammatory cytokines, and calcium signaling [44, 45]. In PMNs MyD88, a downstream signaling adapter protein of TLR signaling, was shown to be required for mitochondrial ROS production indicating a role for TLR signaling in PMNs as well [26]. The previously reported increase in mitochondrial ROS by A2BR^-/-^ PMNs infected with pneumococci opsonized with sera from naïve hosts are most likely complement mediated. While C3b is the component of complement that acts as opsonin additional complement components have been shown to effect mitochondrial responses in other cell types [27]. Complement receptor C5a was found to be expressed internally on mitochondria in monocytes and activation of this receptor on mitochondria by C5a induced mitochondrial ROS production in a dose dependent manner [46]. Complement receptor C3a was also found to localize to the mitochondria and inhibit respiratory function in human retinal pigment epithelial cells [47]. Overall, these findings suggest that additional non-ROS related mechanisms of intracellular pneumococcal killing are being regulated by A2B receptor signaling in young mice.

Fusion of primary and secondary neutrophil granules with the phagosome is necessary for efficient intracellular microbial killing following phagocytosis [38, 48]. CRAMP and the human homologue LL-37 is an important mediator of pneumococcal killing. We found that intracellular levels of secondary granular component CRAMP were elevated in PMNs from A2BR^-/-^ mice and infection of these PMNs with *S. pneumoniae* opsonized with HI Immune sera significantly increased intracellular CRAMP compared to uninfected controls. CRAMP deficient mice were found to have higher bacterial burden and mortality during pneumococcal meningitis [49]. Aged mice were found to have prolonged nasal colonization of *S. pneumoniae* which was associated with diminished CRAMP expression [50]. Additionally aged mice had decreased intracellular killing of pneumococcus following antibody mediated uptake and decreased levels of intracellular CRAMP [11]. In humans, LL-37 was found to be effective against both clinical and laboratory strains of *S. pneumoniae* [51]. Therefore, increased intracellular CRAMP levels could account for increased intracellular killing following antibody mediated uptake in the absence of A2BR signaling.

Neutrophils are necessary for protection against *S. pneumoniae* in a vaccinated host as depletion of PMNs in vaccinated young mice impaired resistance to infection [11]. This study identified EAD signaling through the A2B receptor as a pathway that impairs PMN function in young, vaccinated mice. Extracellular adenosine is a damage associated molecular pattern (DAMP) as it is released upon cellular damage which can be due to infection and inflammation. Cellular damage significantly raises adenosine levels in the extracellular environment from baseline and signaling through EAD receptors occurs [19]. A2BR is a low affinity receptor and is activated when adenosine levels are high. Since adenosine levels rise as infection and inflammation increase it is possible that A2BR signaling acts as a feedback signal to limit PMN activity as ROS, NETs, and antimicrobial peptides can contribute to host cellular damage [52].

Following pneumococcal infection extracellular adenosine levels in the circulation are higher in unvaccinated aged mice when compared to young controls [20]. Since high levels of EAD activate A2BR, which has been shown to be inhibitory to PMN function, we hypothesized that inhibition of A2BR in vaccinated aged hosts would rescue host susceptibility to disease. However, we found here that inhibition of A2BR signaling was not sufficient to reverse the age-related decline in PMN function or the age-related decline in vaccine protection against pneumococcal infection. In contrast, our prior studies found that activating A1R signaling was sufficient to improve PMN responses to pneumococcal challenge and overall host resistance against pneumococcal infection in aged naïve hosts [20, 21]. Our findings here suggest that EAD signaling does not regulate host responses the same way in aged naive versus vaccinated hosts.

The reasons why targeting A2BR was not sufficient to reverse age-driven defects in PMN responses to antibody bound bacteria are not clear. Since basal A2BR expression is similar on PMNs isolated from young and aged mice it is possible that A2BR function is impaired with age to begin with, so further inhibition has no effect [21]. Alternatively, if A2BR activation is similar in young and aged hosts, it is possible that inflammaging (chronic inflammation that occurs with aging) may affect PMN responses despite EAD signaling occurring. Inflammaging results in increased production of proinflammatory cytokines at baseline, which contributes to immunosenescence and age-related defects in PMN function, as well as release of immature PMNs from the bone marrow [53–55]. While proinflammatory cytokines can stimulate PMN activity, prolonged stimulation of PMNs can make them less responsive upon interaction with a second stimulus, such as bacteria [56, 57]. In support of the role of role for chronic inflammation in impairment of PMN responses, PMNs isolated from patients with the chronic inflammatory condition psoriatic arthritis had diminished ROS production, phagocytosis, NETosis, and MPO release when stimulated with TNF compared to healthy controls [55]. Inhibition of A2B receptor signaling may therefore not be enough to overcome the age-related dysregulation of PMN responses induced by inflammaging. Finally, a possible explanation for why inhibition of A2BR signaling cannot boost resistance of vaccinated aged hosts to infection is the decline in antibody response that occur with age. Following both pneumococcal infection and vaccination, antibodies produced by aged hosts have been shown to have reduced opsonophagocytic capacity [9, 58]. This change in antibody functionality may result in changes in PMN activation with age, reducing the antibodies interacting with phagocytic receptors on the cell surface. Adenosine signaling has been shown to regulate antibody production in response to the pneumococcal polysaccharide vaccine where CD73^-/-^ and A2AR^-/-^ mice had delayed isotype switching to IgG3 [36, 59]. This suggests that impaired adenosine signaling with age may contribute to worsening vaccine responses and inhibition of A2BR *in vivo* may not be able to overcome these age-related defects. In summary, this study has identified a targetable pathway that can boost immune responses in young, PCV vaccinated mice. This work expands upon the signaling pathways that control vaccine-mediated responses across host age.

## AUTHOR CONTRIBUTIONS

SRS designed research, conducted research, analyzed data, and wrote paper. APL and MCB conducted research and analyzed data. ENBG designed research, wrote the paper, and had responsibility for final content. All authors read and approved the final manuscript.

## DATA AVAILABILITY STATEMENT

The raw data supporting the conclusions of this article will be made available by the authors on request.

## REFERENCES

1. Grudzinska, F.S., et al., Neutrophils in community-acquired pneumonia: parallels in dysfunction at the extremes of age. Thorax, 2020. 75(2): p. 164–171.

2. Jain, S., et al., Community-Acquired Pneumonia Requiring Hospitalization among U.S. Adults. N Engl J Med, 2015. 373(5): p. 415–27.

3. CDC. Pneumococcal Disease Surveillance and Trends. 2024 September 9, 2024 [cited 2025 February 26]; Available from: https://www.cdc.gov/pneumococcal/php/surveillance/index.html.

4. Shenoy, A.T. and C.J. Orihuela, Anatomical site-specific contributions of pneumococcal virulence determinants. Pneumonia (Nathan), 2016. 8.

5. CDC, Centers for Disease Control and Prevention, Active Bacterial Core Surveillance (ABCs) Report Emerging Infections Network Streptococcus pneumoniae, 2022. 2022.

6. Miwako Kobayashi, A.J.L., Ryan Gierke, Wei Xing, Emma Accorsi, Pedro Moro, Mini Kamboj, George A. Kuchel, Robert Schnechter, Jamie Loehr, Adam L. Cohen, Expanded Recommendations for Use of Pneumococcal Conjugate Vaccine Among Adults Aged ≥ 50 Years: Recommendations of the Advisory Committee on Immunization Practices— United States, 2024. MMWR Morbidity and Mortality Weekly Report, 2025. 74(1);1–8.

7. Bonten, M.J., et al., Polysaccharide conjugate vaccine against pneumococcal pneumonia in adults. N Engl J Med, 2015. 372(12): p. 1114–25.

8. Suzuki, M., et al., Serotype-specific effectiveness of 23-valent pneumococcal polysaccharide vaccine against pneumococcal pneumonia in adults aged 65 years or older: a multicentre, prospective, test-negative design study. Lancet Infect Dis, 2017. 17(3): p. 313–321.

9. Simell, B., et al., Aging reduces the functionality of anti-pneumococcal antibodies and the killing of Streptococcus pneumoniae by neutrophil phagocytosis. Vaccine, 2011. 29(10): p. 1929–34.

10. Gingerich, A.D. and J.J. Mousa, Diverse Mechanisms of Protective Anti-Pneumococcal Antibodies. Front Cell Infect Microbiol, 2022. 12: p. 824788.

11. Simmons, S.R., et al., The Age-Driven Decline in Neutrophil Function Contributes to the Reduced Efficacy of the Pneumococcal Conjugate Vaccine in Old Hosts. Frontiers in Cellular and Infection Microbiology, 2022. 12.

12. Romero-Steiner, S., et al., Reduction in functional antibody activity against Streptococcus pneumoniae in vaccinated elderly individuals highly correlates with decreased IgG antibody avidity. Clin Infect Dis, 1999. 29(2): p. 281–8.

13. Simmons, S.R., et al., Older but Not Wiser: the Age-Driven Changes in Neutrophil Responses during Pulmonary Infections. Infect Immun, 2021. 89(4).

14. Pechous, R.D., With Friends Like These: The Complex Role of Neutrophils in the Progression of Severe Pneumonia. Front Cell Infect Microbiol, 2017. 7: p. 160.

15. Kolaczkowska, E. and P. Kubes, Neutrophil recruitment and function in health and inflammation. Nat Rev Immunol, 2013. 13(3): p. 159–75.

16. Rolston, K.V., The spectrum of pulmonary infections in cancer patients. Curr Opin Oncol, 2001. 13(4): p. 218–23.

17. Garvy, B.A. and A.G. Harmsen, The importance of neutrophils in resistance to pneumococcal pneumonia in adult and neonatal mice. Inflammation, 1996. 20(5): p. 499–512.

18. Haskó, G. and B.N. Cronstein, Adenosine: an endogenous regulator of innate immunity. Trends Immunol, 2004. 25(1): p. 33–9.

19. Barletta, K.E., K. Ley, and B. Mehrad, Regulation of neutrophil function by adenosine. Arterioscler Thromb Vasc Biol, 2012. 32(4): p. 856–64.

20. Simmons, S.R., et al., Activating A1 adenosine receptor signaling boosts early pulmonary neutrophil recruitment in aged mice in response to Streptococcus pneumoniae infection. Immun Ageing, 2024. 21(1): p. 34.

21. Bhalla, M., et al., Extracellular adenosine signaling reverses the age-driven decline in the ability of neutrophils to kill Streptococcus pneumoniae. Aging Cell, 2020: p. e13218.

22. van der Hoeven, D., E.T. Gizewski, and J.A. Auchampach, Activation of the A(3) adenosine receptor inhibits fMLP-induced Rac activation in mouse bone marrow neutrophils. Biochem Pharmacol, 2010. 79(11): p. 1667–73.

23. Ernens, I., et al., Adenosine inhibits matrix metalloproteinase-9 secretion by neutrophils: implication of A2a receptor and cAMP/PKA/Ca2+ pathway. Circ Res, 2006. 99(6): p. 590–7.

24. Barletta, K.E., et al., Adenosine A(2B) receptor deficiency promotes host defenses against gram-negative bacterial pneumonia. Am J Respir Crit Care Med, 2012. 186(10):p. 1044–50.

25. Bou Ghanem, E.N., et al., Extracellular Adenosine Protects against Streptococcus pneumoniae Lung Infection by Regulating Pulmonary Neutrophil Recruitment. PLoS Pathog, 2015. 11(8): p. e1005126.

26. Herring, S.E., et al., Mitochondrial ROS production by neutrophils is required for host antimicrobial function against Streptococcus pneumoniae and is controlled by A2B adenosine receptor signaling. PLoS Pathog, 2022. 18(11): p. e1010700.

27. Vandendriessche, S., et al., Complement Receptors and Their Role in Leukocyte Recruitment and Phagocytosis. Front Cell Dev Biol, 2021. 9: p. 624025.

28. Vitharsson, G., et al., Opsonization and antibodies to capsular and cell wall polysaccharides of Streptococcus pneumoniae. J Infect Dis, 1994. 170(3): p. 592–9.

29. García-García, E. and C. Rosales, Signal transduction during Fc receptor-mediated phagocytosis. J Leukoc Biol, 2002. 72(6): p. 1092–108.

30. Lim, J. and N.A. Hotchin, Signalling mechanisms of the leukocyte integrin αMβ2: current and future perspectives. Biol Cell, 2012. 104(11): p. 631–40.

31. Kobayashi, S.D., et al., Global changes in gene expression by human polymorphonuclear leukocytes during receptor-mediated phagocytosis: cell fate is regulated at the level of gene expression. Proc Natl Acad Sci U S A, 2002. 99(10): p. 6901–6.

32. Greene, N.G., et al., Peptidoglycan Branched Stem Peptides Contribute to Streptococcus pneumoniae Virulence by Inhibiting Pneumolysin Release. PLoS Pathog, 2015. 11(6): p. e1004996.

33. Siwapornchai, N., et al., Extracellular adenosine enhances the ability of PMNs to kill Streptococcus pneumoniae by inhibiting IL-10 production. J Leukoc Biol, 2020.

34. Swamydas, M. and M.S. Lionakis, Isolation, purification and labeling of mouse bone marrow neutrophils for functional studies and adoptive transfer experiments. J Vis Exp, 2013(77): p. e50586.

35. Fante, M.A., et al., Heat-Inactivation of Human Serum Destroys C1 Inhibitor, Pro-motes Immune Complex Formation, and Improves Human T Cell Function. Int J Mol Sci, 2021. 22(5).

36. Allard, D., et al., CD73-A2a adenosine receptor axis promotes innate B cell antibody responses to pneumococcal polysaccharide vaccination. PLoS One, 2018. 13(1): p. e0191973.

37. Yin, C. and B. Heit, Armed for destruction: formation, function and trafficking of neutrophil granules. Cell Tissue Res, 2018. 371(3): p. 455–471.

38. Nordenfelt, P. and H. Tapper, Phagosome dynamics during phagocytosis by neutrophils. J Leukoc Biol, 2011. 90(2): p. 271–84.

39. Sun, Y. and P. Huang, Adenosine A2B Receptor: From Cell Biology to Human Diseases. Front Chem, 2016. 4: p. 37.

40. Antonioli, L., et al., Adenosine signaling and the immune system: When a lot could be too much. Immunol Lett, 2019. 205: p. 9–15.

41. Junger, W.G., Immune cell regulation by autocrine purinergic signalling. Nat Rev Immunol, 2011. 11(3): p. 201–12.

42. Fredholm, B.B., et al., Comparison of the potency of adenosine as an agonist at human adenosine receptors expressed in Chinese hamster ovary cells. Biochem Pharmacol, 2001. 61(4): p. 443–8.

43. Nguyen, G.T., E.R. Green, and J. Mecsas, Neutrophils to the ROScue: Mechanisms of NADPH Oxidase Activation and Bacterial Resistance. Front Cell Infect Microbiol, 2017. 7: p. 373.

44. Zorov, D.B., M. Juhaszova, and S.J. Sollott, Mitochondrial reactive oxygen species (ROS) and ROS-induced ROS release. Physiol Rev, 2014. 94(3): p. 909–50.

45. Shekhova, E., Mitochondrial reactive oxygen species as major effectors of antimicrobial immunity. PLoS Pathog, 2020. 16(5): p. e1008470.

46. Niyonzima, N., et al., Mitochondrial C5aR1 activity in macrophages controls IL-1β production underlying sterile inflammation. Sci Immunol, 2021. 6(66): p. eabf2489.

47. Ishii, M., et al., Mitochondrial C3a Receptor Activation in Oxidatively Stressed Epithelial Cells Reduces Mitochondrial Respiration and Metabolism. Front Immunol, 2021. 12: p. 628062.

48. Borregaard, N., O.E. Sørensen, and K. Theilgaard-Mönch, Neutrophil granules: a library of innate immunity proteins. Trends Immunol, 2007. 28(8): p. 340–5.

49. Merres, J., et al., Role of the cathelicidin-related antimicrobial peptide in inflammation and mortality in a mouse model of bacterial meningitis. J Innate Immun, 2014. 6(2): p. 205–18.

50. Krone, C.L., et al., Impaired innate mucosal immunity in aged mice permits prolonged Streptococcus pneumoniae colonization. Infect Immun, 2013. 81(12): p. 4615–25.

51. Sharma, P., et al., Differential Expression of Antimicrobial Peptides in Streptococcus pneumoniae Keratitis and STAT3-Dependent Expression of LL-37 by Streptococcus pneumoniae in Human Corneal Epithelial Cells. Pathogens, 2019. 8(1).

52. Weiss, S.J., Tissue destruction by neutrophils. N Engl J Med, 1989. 320(6): p. 365–76.

53. López-Otín, C., et al., Hallmarks of aging: An expanding universe. Cell, 2023. 186(2): p.243–278.

54. Li, X., et al., Inflammation and aging: signaling pathways and intervention therapies. Signal Transduct Target Ther, 2023. 8(1): p. 239.

55. Leliefeld, P.H., et al., The role of neutrophils in immune dysfunction during severe inflammation. Crit Care, 2016. 20: p. 73.

56. Martin, A., et al., Effect of vitamin E on hydrogen peroxide production by human vascular endothelial cells after hypoxia/reoxygenation. Free Radic Biol Med, 1996. 20(1): p. 99–105.

57. Divangahi, M., et al., Trained immunity, tolerance, priming and differentiation: distinct immunological processes. Nat Immunol, 2021. 22(1): p. 2–6.

58. Schenkein, J.G., S. Park, and M.H. Nahm, Pneumococcal vaccination in older adults induces antibodies with low opsonic capacity and reduced antibody potency. Vaccine, 2008. 26(43): p. 5521–6.

59. Schmiel, S.E., et al., Cutting Edge: Adenosine A2a Receptor Signals Inhibit Germinal Center T Follicular Helper Cell Differentiation during the Primary Response to Vaccination. J Immunol, 2017. 198(2): p. 623–628.

