## Supplemental Figure 1 for "Adenosine 2B receptor signaling impairs vaccine-mediated protection against pneumococcal infection in young hosts by blunting neutrophil killing of antibody opsonized bacteria"

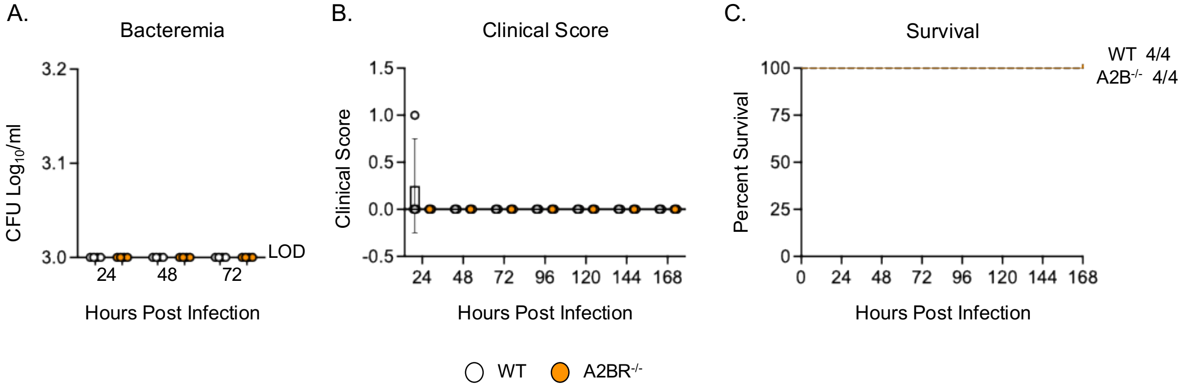


**Supplemental Figure 1:** Young (2-3 months) WT C57BL/6 and A2BR$^{-/-}$ were vaccinated with PCV and 4 weeks later infected i.t with $2x10^6$ CFU of *S. pneumoniae* TIGR4. At 25, 48, and 72 hpi blood was collected and plated for CFU to assess bacteremia (A). Mice were also monitored for 7 days and assessed for clinical signs of disease (B) and survival (C). Pooled data from n=4 mice per group are shown.
